# A comprehensive human minimal gut metagenome extends the host’s metabolic potential

**DOI:** 10.1101/818138

**Authors:** Marcos Parras-Moltó, Daniel Aguirre de Cárcer

**Affiliations:** Departamento de Biología, Universidad Autónoma de Madrid, Madrid, 28049 Spain.

**Keywords:** Human gut Microbiome, 16S rRNA gene, PICRUSTs, Community Assembly, Metagenomics

## Abstract

Accumulating evidence suggests that humans should be considered as holobionts in which the gut microbiota plays essential functions. Initial metagenomic studies reported a pattern of shared genes in the gut microbiome of different individuals, leading to the definition of the minimal gut metagenome as the set of microbial genes necessary for homeostasis, and present in all healthy individuals. Despite its interest, this concept has received little attention following its initial description in terms of various ubiquitous pathways in Western cohorts. This study analyzes the minimal gut metagenome of the most comprehensive dataset available, including individuals from agriculturalist and industrialist societies, also embodying highly diverse ethnic and geographical backgrounds. The outcome, based on metagenomic predictions for community composition data, resulted in a minimal metagenome comprising 3,412 gene clusters, mapping to 1,856 reactions and 128 metabolic pathways predicted to occur across all individuals. These results were substantiated by the analysis of two additional datasets describing the microbial community compositions of larger Western cohorts, as well as a substantial shotgun metagenomics dataset. Subsequent analyses showed the plausible metabolic complementarity provided by the minimal gut metagenome to the human genome.

---

The study of the human gut microbiome has drawn from different disciplines (e.g. microbiology, ecology, genomics), and has substantiated the idea that humans should be considered as holobionts (1) in which the gut microbiota plays essential functions (2, 3). Knowledge of what constitutes a healthy gut microbiome is regarded as pivotal (4) for the development of predictive models for diagnosis and management of gut microbiome-related maladies. However, the strong inter-subject variability in community composition observed in cross-sectional studies (5) hindered an early definition of a set of bacterial species common to all healthy humans (6). While, recent efforts have been able to detect such a health-related set in terms of shared taxonomic assignments (4, 7), and more precisely in terms of shared 16S sequence clusters of varying phylogenetic depth (8), the idea that a healthy gut microbiome ‘core’ may exist only in terms of function (9) remains widespread.

In this regard, early high-throughput shotgun metagenomic studies already reported a strong pattern of shared genes in the gut microbiome of different individuals (10, 11). These results led to the definition of a novel concept; the minimal gut metagenome (11), defined as the set of microbial genes necessary for the homeostasis of the whole gut ecosystem, and expected to be present in all healthy humans. The idea that the gut microbiome provides a specific set of functionalities shared by all individuals is intuitive. However, it is still unclear whether these functionalities could arise from a shared set of genes or from different combinations of genes. Moreover, if the host were to play a greatly diminished role as a selective force on its resident gut microbiome, when compared to external factors such as diet, then there would be no set of microbial functionalities shared by all humans.

Nevertheless, despite its potential as a conceptual framework with which to study the gut ecosystem, the minimal gut metagenome concept has received little attention in the literature following its initial definition and description in terms of various ubiquitous metabolic pathways (9-11) and recent description of prevalent pathways in a larger cohort (12).

Hence, the aim of the present study is to recapitulate the minimal human gut metagenome conceptual framework, and provide a proof-of-concept of its utility. More specifically, we set out to identify the ‘core genes’ (defined as the set of genes detected in all individuals), jointly comprising the minimal gut metagenome, as well as the ‘core reactions’ (defined as the set of metabolic reactions detected in all individuals). According to the minimal gut metagenome concept, the former should be related to gut homeostasis at large (i.e. not only metabolic homeostasis). On the other hand, knowledge on the latter should improve our understanding of the gut microbiome’s ability to augment human metabolism.

For knowledge of the minimal gut metagenome to be most useful, it should pertain more to *Homo sapiens* as a species, and hence should not be solely focused on Western cohorts. Unfortunately, most human gut shotgun metagenomic datasets are very restricted in terms of lifestyles and ethnicities, mostly arising from Western and(or) industrialist cohorts (9-13).

In this study, 16S rRNA gene-based metagenomic predictions were employed in the assessment of the minimal human gut metagenome to be able to profit from the more comprehensive 16S datasets. These datasets greatly outclass available human gut shotgun metagenomic datasets in terms of cohort size, geographic distribution, ethnic and lifestyle diversity, and to a certain extent depth of sequencing. In a sense, one read in a shotgun metagenomics dataset represents one gene count, while one read in a 16S amplicon survey represents, *via* metagenomic prediction, one genome count. However, the use of metagenomic predictions presents various limitations and possible biases, which have been explored previously (14), the most noteworthy being that it only infers the bacterial and archaeal component of the metagenome, is significantly affected by both the quality of available genome annotations and the fact that available genomes are not evenly distributed across the phylogeny, or the lack of perfect one-to-one mapping between genomes and even full-length 16S sequences. Nevertheless, the ability to count almost three orders of magnitude more genes in a metagenomic sample per sequence (with the number of bacterial genes per genome normally in the very few thousand), even as a prediction, is still useful. In this study, functional predictions based on 16S phylogenetic marker gene sequences were obtained using PICRUSt, a computational approach which has shown large and significant correlation in predicting metagenomic abundances from 16S measurements (Spearman r = 0.82, p < 0.001) and synthetic communities (Spearman r = 0.9, p < 0.001)(14). To date, PICRUSt has been used in a myriad of scientific works and different research scenarios, such as the analysis of environmental samples (15), medically-relevant communities (16), or *in vitro* assemblies (17).This study analyzes the minimal gut metagenome of the most comprehensive dataset available (dataset *Global*: 382 individuals from rural Malawi, metropolitan U.S.A., and Venezuelan Amerindians(18). See **Table 1**)), which, despite its comparatively smaller cohort size, is far more inclusive in terms of global distribution, lifestyle, and ethnicity, specifically including agriculturalist, and industrialist societies from three continents.

We compare the Global dataset with two larger Western cohorts (dataset *Flemish*: 873 individuals from Belgium (4); and dataset *Twins: 2,727 individuals from U.K.* (19)), as well as to a substantial shotgun metagenomics dataset (Dataset *Shotgun*: KEGG Orthology identifiers (KOs) (20) abundances from 123 individuals from U.S.A., Europe, and China. Obtained from Bradley and Pollard 2017 (21)), and compared with the human genome to assess the degree to which the minimal metagenome may complement and expand its host’s metabolic potential.

## RESULTS

The authors of the PICRUSt paper state that there is a significant negative correlation (Spearman r = −0.4, P < 0.001) between NSTI values and Spearman correlation between empirical shotgun metagenome abundances and PICRUSt predictions based on 16S sequences.(14). Here, NSTI values for the different sample sets of *Global* (0.135±0.021, 0.098±0.018, and 0.131±0.023 for Malawian, U.S.A., and Venezuelan samples, respectively; see **Suppl. Fig 1**) were lower (generally correlated with higher correlation between metagenomic measurements and 16S predictions) than those previously reported for soil samples (0.17±0.02) which showed a significant [P<0.001] correlation between predictions and matched shotgun metagenomics assignments (14). Also, the more extreme NSTI values reported for the Human Microbiome Project dataset, with NSTI values ranging 0.10-0.15, still presented high correlation coefficients between metagenomic measurements and 16S predictions (14).

The results show that 5,865 KO groups were predicted as present in *Global*’s pan-metagenome, while the minimal gut metagenome represented 3,412 KOs (i.e. core genes), which can in turn be mapped to 1,856 reactions (i.e. core reactions) and 128 complete metabolic pathways (**Additional file 1**).

As could be expected, lowering the prevalence threshold used to define core reactions (100%) increased the number of core reactions, but mainly in a gentle-slope linear fashion (**Suppl. Fig. 2**). The core metagenome was very similar among the three distinct sample sets comprising *Global* (**Figure 1A**), with U.S.A.’s set showing the smallest set of core reactions, and less overlap with Malawian and Venezuelan samples. On the other hand, *Global*’s core reaction set was comparatively similar to those obtained using Western-like datasets *Twins* and *Flemish* (**Figure 1B**).

**Figure 1.**
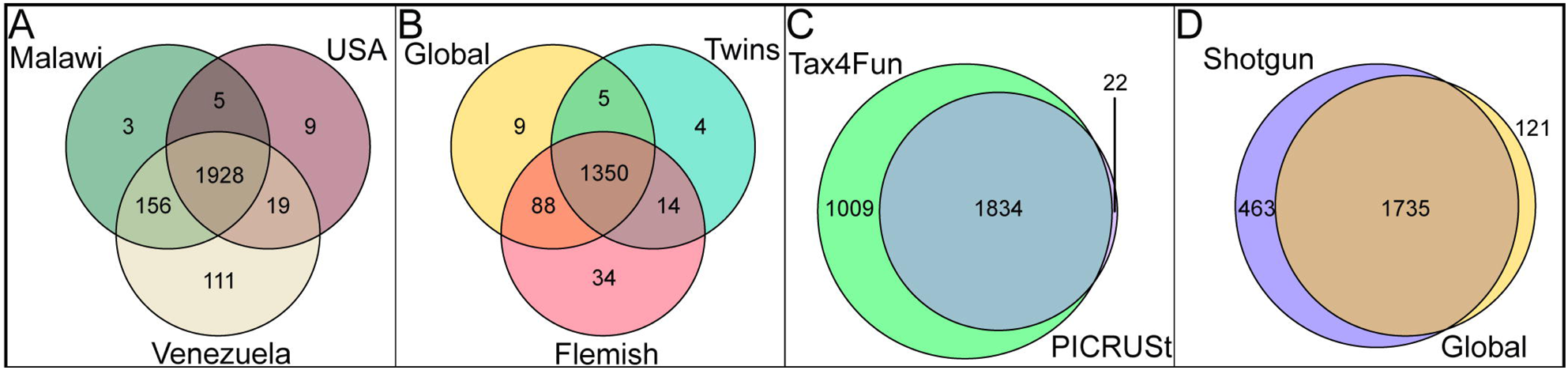
Venn diagrams depicting the overlap in core reactions between different datasets and software. **Panel A**: Different sample sets within *Global*. Values refer to the analysis with the same number of individuals per population (50). **Panel B**: Different 16S datasets. Values refer to the analysis with the same number of sequences per sample (8,000). **Panel C**: Differences between metagenomic prediction software. **Panel D**: Differences between 16S (*Global*) and shotgun metagenomics (*Shotgun*) datasets.

The presented core reactions were predicted from 16S profiles using an ancestral-state reconstruction algorithm (PICRUSt). However, the set of core reactions was substantiated by the use of Tax4fun (22), a taxonomy assignments-based approach (**Figure 1C**). PICRUSt’s predictions seem conservative (more appropriate for a minimum estimate, as intended) since they are a subset of Tax4fun predictions. More importantly, *Global*’s core reaction set presented a high overlap to that obtained from a substantial shotgun metagenomics dataset targeting the human gut microbiome (21), chosen among those publicly available based on the number of individuals and geographic and ethnic distribution (**Figure 1D**). The 463 reactions described as core in *Shotgun* but not in *Global* (**Figure 1D**) likely arise from the smaller size of the *Shotgun*’s cohort as well as its increased lifestyle, environmental and genetic homogeneity (**Table 1**). On the other hand, the great majority of core reactions in *Global* not described as core in *Shotgun* still presented a very high prevalence in the dataset (**Suppl. Fig. 3**); 1,735 out of 1,856 (93.5%) core reactions in *Global* are also core reactions (100% prevalence) in *Shotgun*. Only 37 (2%) core reactions in *Global* have a prevalence level < 95% in *Shotgun*, and 6 (0.32%) reactions have a prevalence level below 75%. No apparent shared functional or taxonomic origin affiliation was found for these six reactions. Within the *Global* dataset, there was a positive correlation between prevalence and average abundance (**Suppl. Fig. 4**). Nevertheless, while all core reactions featured relatively high average abundance values, many similarly abundant reactions presented lower prevalence values.

**Table 1.** Datasets’ characteristics

| **Name** | **Geographic distribution** | **Number of individuals** | **Sequence depth<sup>1</sup>** | **Read length<sup>2</sup>** | **Sequencing technology<sup>3</sup>** |
| --- | --- | --- | --- | --- | --- |
| **Global** | Malawi, USA, Venezuela | 382 | >300K | 100 | GAIIx |
| **Twins** | UK | 2,727 | >15K | 2x250 | MiSeq |
| **Flemish** | Belgium | 873 | >8K | 2x250 | MiSeq |
| **Shotgun** | USA, Europe, China | 123 | 15M | 2x75, 2x100 | GAIIx, HiSeq |

<sup>1</sup> Values represent final sequence depth per sample before analysis (i.e. after chimera removal and subsampling to common depth). <sup>2</sup> in bp. <sup>3</sup> Illumina

In addition to providing an improved description of the human minimal gut metagenome, the present study aimed at assessing its complementarity to the human genome. In this regard, the metabolic complementarity judged by the Metabolic Complementarity Index (23) was >2 times larger when considering the human metabolism being complemented by *Global*’s minimal gut metagenome, when compared to the inverse (0.0807 and 0.0386, respectively).

Considering two metabolites as linked if they represent the substrate and product of a core reaction, within the overall metabolic map (**Figure 2**, **Suppl. Fig. 5**) 199 microbial metabolites link with 89 *Homo sapien*s metabolites through 256 core reactions, representing the predicted extended metabolic capability of the human holobiont provided by its gut ecosystem. Additionally, the map pinpoints 55 core reactions and 84 metabolites with no apparent connection to *Homo sapiens* metabolism, as well as 36 core reactions able to link *Homo sapiens* metabolites by reactions different to those carried-out by enzymes encoded within the human genome.

**Figure 2.**
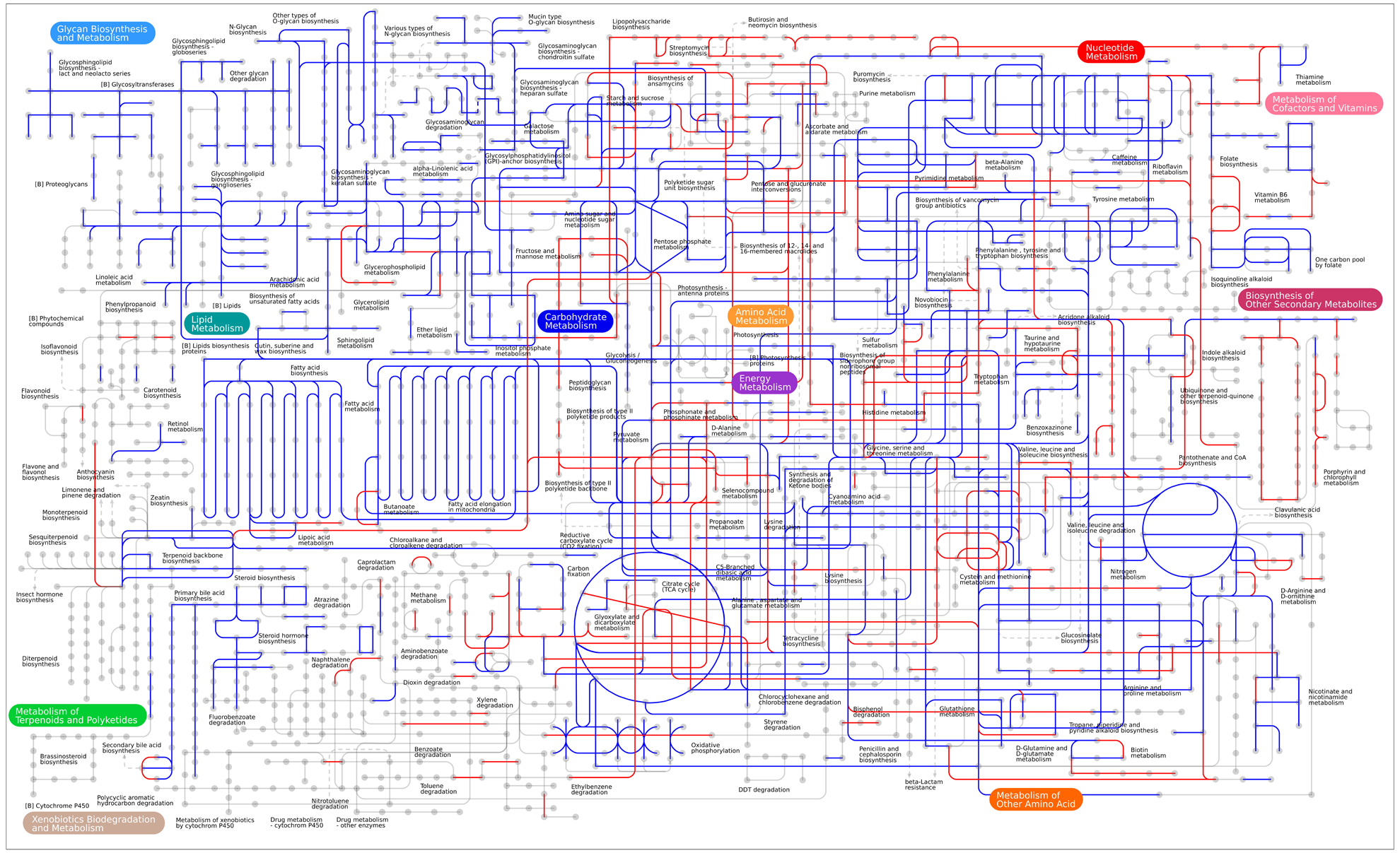
The minimal gut metagenome extends human metabolic potential. Nodes in the map correspond to chemical compounds and edges represent enzymatic reactions. The figure provides an iPath2.0 representation of KEGG metabolic pathways, where reactions catalyzed by enzymes encoded in the human genome appear in blue, while core reactions of the human gut pan-microbiome not encoded also by the human genome, appear in red.

Not surprisingly, several core reactions are implicated in the production of short-chain fatty acids (SCFAs), such as butyrate and acetate, which are known to have an active role in normal human physiology (e.g. fuel for several cell types, regulation of gene expression, differentiation, and inflammation) (24, 25). Another hallmark of the predicted minimal gut metagenome relates to the presence of core reactions implicated in the production of several vitamins (B1, B2, B5, B6, B9, H, K1, K2, L1, coenzyme B12), several of which had previously been shown to be produced by common gut commensals (26).

## DISCUSSION

The NSTI values that we obtained for human gut microbiome samples fall within the range of NSTI values for samples in the PICRUSt validation that had high correlation between metagenomic abundance measurements and 16S predictions.(14). In this regard, an enhanced and updated report on the utility, correlation between predicted and experimental measurements, and accuracy of PICRUSt’s predictions would be welcomed by the community, more so since this area of development seems to remain active (27, 28). The values obtained were not homogenous among the three distinct sample sets in *Global*, with values for both the Venezuelan and Malawian samples being roughly 35% higher than that of the U.S.A. samples. In this regard, the detected functional overlap could somewhat be inflated since the reference genome set employed is likely biased towards strains obtained from industrialist countries.

Interestingly, the results indicate that the U.S.A. population restricted the number of detected core reactions, since Venezuela and Malawian samples presented an additional 156 reactions with 100% prevalence in their joint dataset, compared to <20 exclusively shared with 100% prevalence between U.S.A. samples and any of the other groups. Moreover, these values may be conservative, since the reference genomes may be biased towards bacterial strains more frequent in industrialist countries. This reduction in functional overlap provides circumstantial support to the emerging concern that industrialist populations may have lost the microbial diversity needed to adequately sustain a healthy host (29).

The results presented herein are influenced by the fact that the metagenomic prediction approach employed is, to a certain extent, biased, as explained before. As such, the core genes and reactions reported should be taken cautiously. Thus, validation of each particular core reaction in the ecosystem, as well as the possibility of each core metabolite traversing the membrane, along with its potential significance to the host, is beyond the scope of this study. Nevertheless, returning to the three possible scenarios of shared functionality in the human gut pan-microbiome postulated above; i) no shared functionality, ii) shared functionality related to different combinations of genes, and iii) shared functionality related to a shared combination of genes, the results are strongly supportive of the latter. Thus, we believe that the minimal gut metagenome idea indeed represents a potentially useful conceptual framework with which to improve our knowledge of the role played by the human gut microbiome on maintaining host homeostasis.

The results also indicate that the human gut minimal metagenome may extensively contribute to the human holobiont’s metabolic potential. The core reactions reported here represent a highly restrictive set, since reactions need to be present in all subjects to achieve the ‘core’ status. Most importantly, these core reactions were predicted as present in all subjects from a cohort including individuals from agriculturalist and industrialist societies, also embodying highly diverse genetic, ethnic, and geographical backgrounds. Furthermore, the results were validated using additional large-cohort datasets, as well as a substantial shotgun metagenomics dataset. Hence, the described minimal gut metagenome now pertains more to *Homo sapiens* as a species, rather than to industrialist societies of particular ethnic and geographical backgrounds. Finally, our results seem to indicate that the minimal metagenome has a greater role in complementing the human metabolism than the other way around.

## MATERIALS AND METHODS

### Datasets

All datasets comprised 16S rRNA gene sequences obtained using primer pair F515-R806 targeting the V4 hypervariable region, with the exception of dataset *Shotgun* which included KOs abundances obtained through shotgun sequencing of metagenomic DNA (21). All sequence data was derived from stool samples from healthy subjects over 3 years old, with no history of recent antibiotic treatment prior to sampling (see **Table 1**).

### Metagenomic predictions

QIIME (30) scripts were employed during initial sequence processing (Additional file 2). Briefly, datasets were independently processed as follows; first subsampled to the minimum common depth. Then, chimeric sequences were identified with *usearch61* (31) and removed. Finally, sequences were clustered into OTUs using Greengenes (32) 0.97 representative sequence dataset (May 2013) as reference using *usearch61*. Subsequently, PICRUSt scripts were employed to first normalize OTU abundances by 16S rRNA gene copy number, and then transform normalized OTU abundances into KO abundances. Correlation between predictions and measurements was evaluated using NSTI as a proxy for the Spearman coefficient, as they are strongly negatively and significantly correlated (14). Tax4Fun (22), an alternative metagenome prediction pipeline, was also employed with *Global* dataset following the suggested standard procedure.

Since more than one KO group may carry out a particular reaction, KO abundances were mapped to KEGG reactions. In cases where a KO mapped to more than one reaction, all reactions linked to the KO were scored. KOs and reactions appearing in all individuals in the datasets were defined as ‘core’. Finally, the MinPath algorithm (33) was used for biological pathway reconstruction from core KOs.

### Metabolic complementarity assessments

Host-microbiome cooperation was assessed with NetCooperate (23) using the Metabolic Complementarity Index. This index provides a quantification of the extent to which two species may support one another through biosynthetic complementarity. There is no threshold for ‘complementarity’ and ‘no complementarity’, and hence the metrics have to be employed in a comparative manner (23). Here, the index was used to study both moieties of the human holobiont; the human genome and the minimal gut metagenome. Hence, the reciprocal analysis evaluates the relative strength of each moiety complementing the other. To do so, core reactions were transformed into linked KEGG compounds, and then analyzed with NetCooperate. To further assess such complementarity, both the core reactions and the reactions encoded by the human genome were imported into the interactive metabolic pathway explorer iPATH3.0 (34).

## Availability of data and material

The datasets analyzed during the current study are available from their original source (as stated above). Core KOs, Reactions and Compounds are available within **Additional file 1**. Additional intermediate result files and scripts are available from the corresponding author on request for research purposes.

## Authors’ contributions

DA Conceived the idea and wrote the manuscript. MP and DA analyzed the datasets.

## Acknowledgements

We thank the Bioinformatics Unit at CBMSO for their support.

## Additional file 1

Excel file (.xlsx)

Core KOs, Reactions, Compounds and Pathways.

## Additional file 2

Word file (.docx)

QIIME scripts employed.

## FIGURE LEGENDS

**Supplementary Figure 1. Distribution of NSTI values among the three sample sets in *Global***.

**Supplementary Figure 2. The number of core reactions varies with prevalence threshold**. [Linear regression; y=-8.57 + 2899, R^2^=0.92]

**Supplementary Figure 3. Prevalence of *Global* core reactions in *Shotgun***. Dots represent all reactions detected in *Shotgun*. Their prevalence in the dataset is recorded along the y-axis, and those reactions with 100% prevalence in *Global* (core) appear in a different color.

**Supplementary Figure 4. Prevalence Vs. average abundance values in *Global***. Dots represent all reactions predicted in the dataset, core reactions depicted in red.

**Supplementary Figure 5. The gut metagenome extends human metabolic potential.** Nodes in the map correspond to chemical compounds and edges represent enzymatic reactions. The figure provides an iPath2.0 representation of KEGG metabolic pathways, where reactions catalyzed by enzymes encoded in the human genome appear in blue, while reactions of the human gut pan-microbiome not encoded also by the human genome appear in either red (100% prevalence), orange (50% prevalence), or yellow (1% prevalence).

## REFERENCES

1. Bordenstein SR, Theis KR. 2015. Host Biology in Light of the Microbiome: Ten Principles of Holobionts and Hologenomes. PLOS Biology 13:e1002226.

2. Backhed F, Ley RE, Sonnenburg JL, Peterson DA, Gordon JI. 2005. Host-bacterial mutualism in the human intestine. Science 307:1915–1920.

3. Gill SR, Pop M, Deboy RT, Eckburg PB, Turnbaugh PJ, Samuel BS, Gordon JI, Relman DA, Fraser-Liggett CM, Nelson KE. 2006. Metagenomic analysis of the human distal gut microbiome. Science 312:1355–1359.

4. Falony G, Joossens M, Vieira-Silva S, Wang J, Darzi Y, Faust K, Kurilshikov A, Bonder MJ, Valles-Colomer M, Vandeputte D, Tito RY, Chaffron S, Rymenans L, Verspecht C, De Sutter L, Lima-Mendez G, D’hoe K, Jonckheere K, Homola D, Garcia R, Tigchelaar EF, Eeckhaudt L, Fu J, Henckaerts L, Zhernakova A, Wijmenga C, Raes J. 2016. Population-level analysis of gut microbiome variation. Science 352:560–564.

5. Aguirre de Carcer D, Cuiv PO, Wang T, Kang S, Worthley D, Whitehall V, Gordon I, McSweeney C, Leggett B, Morrison M. 2011. Numerical ecology validates a biogeographical distribution and gender-based effect on mucosa-associated bacteria along the human colon. ISME J 5:801–809.

6. Turnbaugh PJ, Ley RE, Hamady M, Fraser-Liggett CM, Knight R, Gordon JI. 2007. The human microbiome project. Nature 449:804–810.

7. Zhang J, Guo Z, Xue Z, Sun Z, Zhang M, Wang L, Wang G, Wang F, Xu J, Cao H, Xu H, Lv Q, Zhong Z, Chen Y, Qimuge S, Menghe B, Zheng Y, Zhao L, Chen W, Zhang H. 2015. A phylo-functional core of gut microbiota in healthy young Chinese cohorts across lifestyles, geography and ethnicities. ISME J 9:1979–1990.

8. Aguirre de Cárcer D. 2018. The human gut pan-microbiome presents a compositional core formed by discrete phylogenetic units. Scientific Reports 8:14069.

9. Turnbaugh PJ, Hamady M, Yatsunenko T, Cantarel BL, Duncan A, Ley RE, Sogin ML, Jones WJ, Roe BA, Affourtit JP, Egholm M, Henrissat B, Heath AC, Knight R, Gordon JI. 2009. A core gut microbiome in obese and lean twins. Nature 457:480–484.

10. Human Microbiome Project Consortium. 2012. Structure, function and diversity of the healthy human microbiome. Nature 486:207–214.

11. Qin J, Li R, Raes J, Arumugam M, Burgdorf KS, Manichanh C, Nielsen T, Pons N, Levenez F, Yamada T, Mende DR, Li J, Xu J, Li S, Li D, Cao J, Wang B, Liang H, Zheng H, Xie Y, Tap J, Lepage P, Bertalan M, Batto JM, Hansen T, Le Paslier D, Linneberg A, Nielsen HB, Pelletier E, Renault P, Sicheritz-Ponten T, Turner K, Zhu H, Yu C, Jian M, Zhou Y, Li Y, Zhang X, Qin N, Yang H, Wang J, Brunak S, Dore J, Guarner F, Kristiansen K, Pedersen O, Parkhill J, Weissenbach J, Bork P, Ehrlich SD. 2010. A human gut microbial gene catalogue established by metagenomic sequencing. Nature 464:59–65.

12. Lloyd-Price J, Mahurkar A, Rahnavard G, Crabtree J, Orvis J, Hall AB, Brady A, Creasy HH, McCracken C, Giglio MG, McDonald D, Franzosa EA, Knight R, White O, Huttenhower C. 2017. Strains, functions and dynamics in the expanded Human Microbiome Project. Nature 550:61.

13. Li J, Jia H, Cai X, Zhong H, Feng Q, Sunagawa S, Arumugam M, Kultima JR, Prifti E, Nielsen T, Juncker AS, Manichanh C, Chen B, Zhang W, Levenez F, Wang J, Xu X, Xiao L, Liang S, Zhang D, Zhang Z, Chen W, Zhao H, Al-Aama JY, Edris S, Yang H, Hansen T, Nielsen HB, Brunak S, Kristiansen K, Guarner F, Pedersen O, Dore J, Ehrlich SD, Bork P. 2014. An integrated catalog of reference genes in the human gut microbiome. Nat Biotechnol 32:834–841.

14. Langille MG, Zaneveld J, Caporaso JG, McDonald D, Knights D, Reyes JA, Clemente JC, Burkepile DE, Vega Thurber RL, Knight R, Beiko RG, Huttenhower C. 2013. Predictive functional profiling of microbial communities using 16S rRNA marker gene sequences. Nat Biotechnol 31:814–821.

15. Bier RL, Voss KA, Bernhardt ES. 2015. Bacterial community responses to a gradient of alkaline mountaintop mine drainage in Central Appalachian streams. ISME J 9:1378–1390.

16. Buffie CG, Bucci V, Stein RR, McKenney PT, Ling L, Gobourne A, No D, Liu H, Kinnebrew M, Viale A, Littmann E, van den Brink MR, Jenq RR, Taur Y, Sander C, Cross JR, Toussaint NC, Xavier JB, Pamer EG. 2015. Precision microbiome reconstitution restores bile acid mediated resistance to Clostridium difficile. Nature 17. 517:205–208.

17. Goldford JE, Lu N, Bajić D, Estrela S, Tikhonov M, Sanchez-Gorostiaga A, Segrè D, Mehta P, Sanchez A. 2018. Emergent simplicity in microbial community assembly. Science 361:469–474.

18. Yatsunenko T, Rey FE, Manary MJ, Trehan I, Dominguez-Bello MG, Contreras M, Magris M, Hidalgo G, Baldassano RN, Anokhin AP, Heath AC, Warner B, Reeder J, Kuczynski J, Caporaso JG, Lozupone CA, Lauber C, Clemente JC, Knights D, Knight R, Gordon JI. 2012. Human gut microbiome viewed across age and geography. Nature 486:222–227.

19. Goodrich JK, Davenport ER, Beaumont M, Jackson MA, Knight R, Ober C, Spector TD, Bell JT, Clark AG, Ley RE. 2016. Genetic Determinants of the Gut Microbiome in UK Twins. Cell Host Microbe 19:731–743.

20. Kanehisa M, Sato Y, Kawashima M, Furumichi M, Tanabe M. 2016. KEGG as a reference resource for gene and protein annotation. Nucleic Acids Res 44:17.

21. Bradley PH, Pollard KS. 2017. Proteobacteria explain significant functional variability in the human gut microbiome. Microbiome 5:017–0244.

22. Asshauer KP, Wemheuer B, Daniel R, Meinicke P. 2015. Tax4Fun: predicting functional profiles from metagenomic 16S rRNA data. Bioinformatics 31:2882–2884.

23. Levy R, Carr R, Kreimer A, Freilich S, Borenstein E. 2015. NetCooperate: a network-based tool for inferring host-microbe and microbe-microbe cooperation. BMC Bioinformatics 16:015–0588.

24. Donohoe DR, Garge N, Zhang X, Sun W, O’Connell TM, Bunger MK, Bultman SJ. 2011. The microbiome and butyrate regulate energy metabolism and autophagy in the mammalian colon. Cell Metab 13:517–526.

25. Furusawa Y, Obata Y, Fukuda S, Endo TA, Nakato G, Takahashi D, Nakanishi Y, Uetake C, Kato K, Kato T, Takahashi M, Fukuda NN, Murakami S, Miyauchi E, Hino S, Atarashi K, Onawa S, Fujimura Y, Lockett T, Clarke JM, Topping DL, Tomita M, Hori S, Ohara O, Morita T, Koseki H, Kikuchi J, Honda K, Hase K, Ohno H. 2013. Commensal microbe-derived butyrate induces the differentiation of colonic regulatory T cells. Nature 504:446–450.

26. Letunic I, Yamada T, Kanehisa M, Bork P. 2008. iPath: interactive exploration of biochemical pathways and networks. Trends Biochem Sci 33:101–103.

27. Douglas GM, Beiko RG, Langille MGI. 2018. Predicting the Functional Potential of the Microbiome from Marker Genes Using PICRUSt. Methods Mol Biol:8728-8723_8711.

28. Douglas GM, Maffei VJ, Zaneveld J, Yurgel SN, Brown JR, Taylor CM, Huttenhower C, Langille MGI. 2019. PICRUSt2: An improved and extensible approach for metagenome inference. bioRxiv:672295.

29. Blaser MJ. 2018. The Past and Future Biology of the Human Microbiome in an Age of Extinctions. Cell 172:1173–1177.

30. Caporaso JG, Kuczynski J, Stombaugh J, Bittinger K, Bushman FD, Costello EK, Fierer N, Pena AG, Goodrich JK, Gordon JI, Huttley GA, Kelley ST, Knights D, Koenig JE, Ley RE, Lozupone CA, McDonald D, Muegge BD, Pirrung M, Reeder J, Sevinsky JR, Turnbaugh PJ, Walters WA, Widmann J, Yatsunenko T, Zaneveld J, Knight R. 2010. QIIME allows analysis of high-throughput community sequencing data. Nat Meth 7:335–336.

31. Edgar RC. 2010. Search and clustering orders of magnitude faster than BLAST. Bioinformatics 26:2460–2461.

32. DeSantis TZ, Hugenholtz P, Larsen N, Rojas M, Brodie EL, Keller K, Huber T, Dalevi D, Hu P, Andersen GL. 2006. Greengenes, a chimera-checked 16S rRNA gene database and workbench compatible with ARB. Appl Environ Microbiol 72:5069–5072.

33. Ye Y, Doak TG. 2009. A parsimony approach to biological pathway reconstruction/inference for genomes and metagenomes. PLoS Comput Biol 5:14.

34. Darzi Y, Letunic I, Bork P, Yamada T. 2018. iPath3.0: interactive pathways explorer v3. Nucleic Acids Res 46:W510–W513.

