## Supplementary figures and images for "A comprehensive human minimal gut metagenome extends the host’s metabolic potential"

### Supplemental Data 1

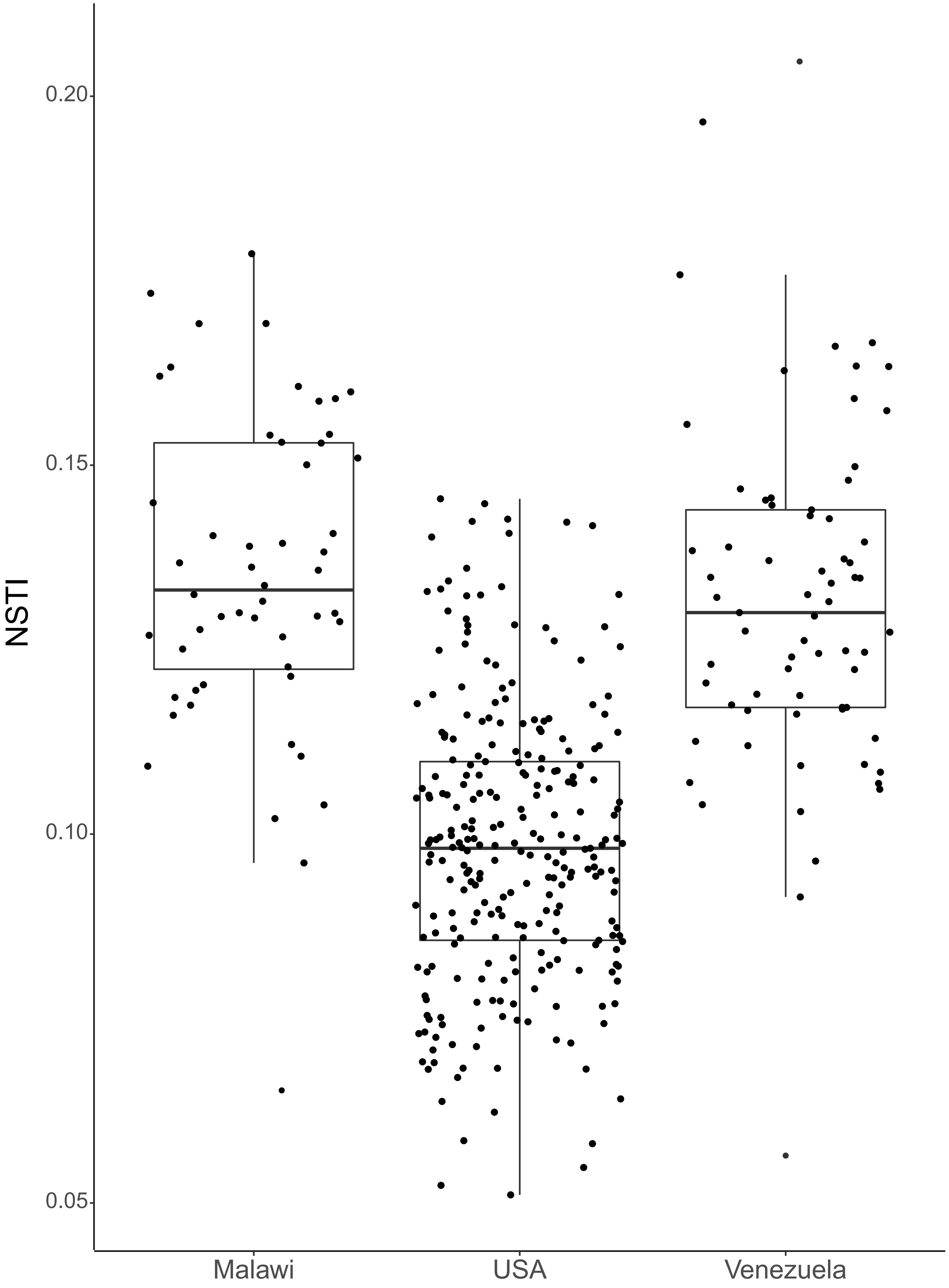

### Supplemental Data 2

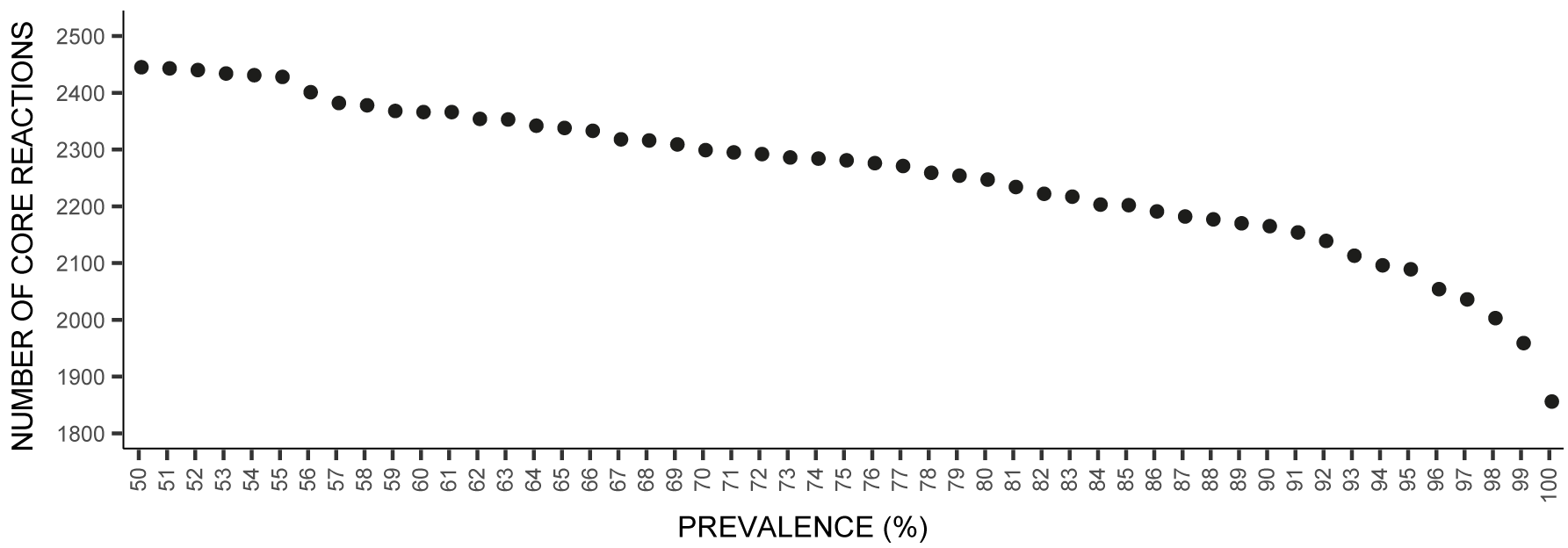

### Supplemental Data 5

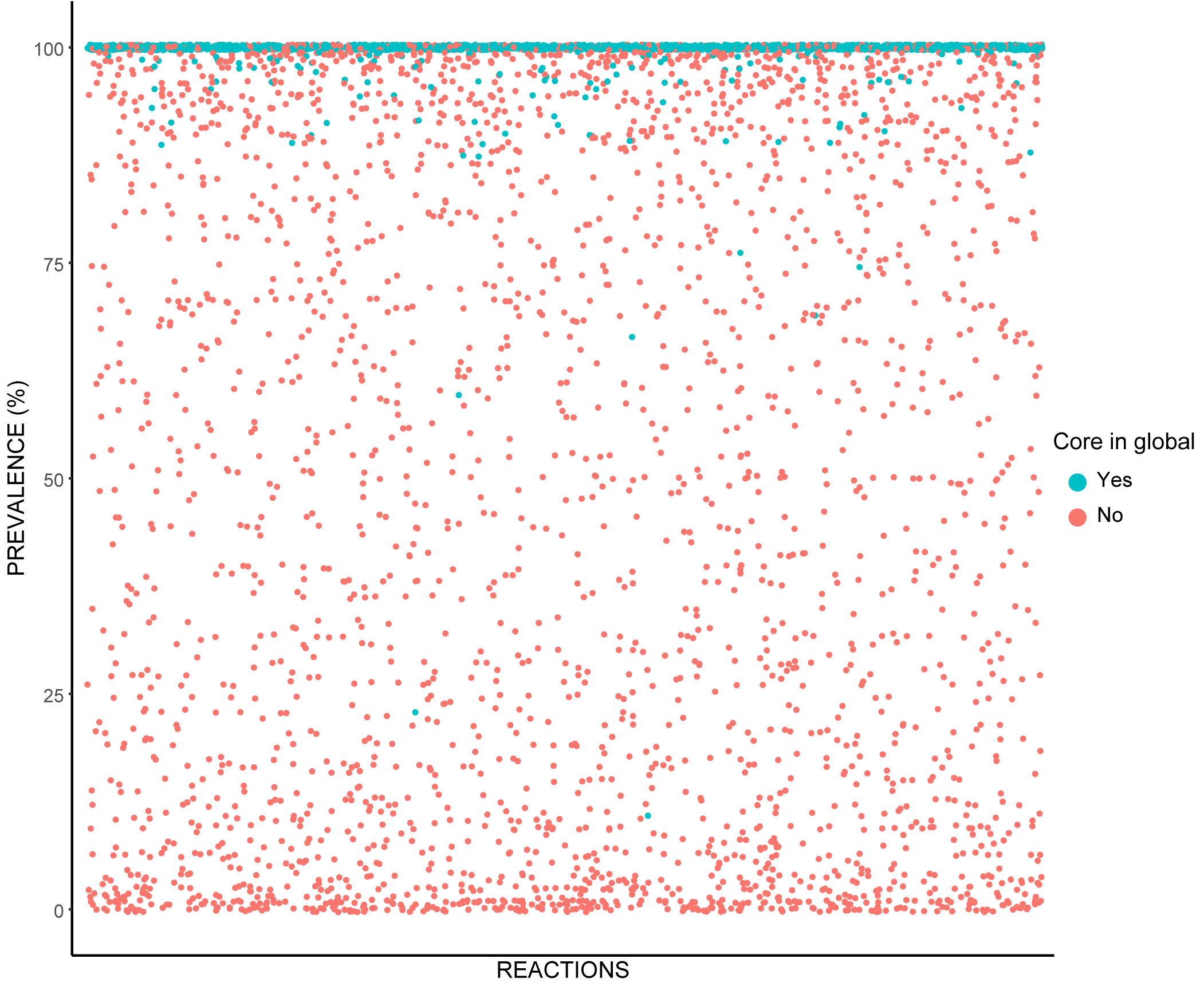

### Supplementary Figure5

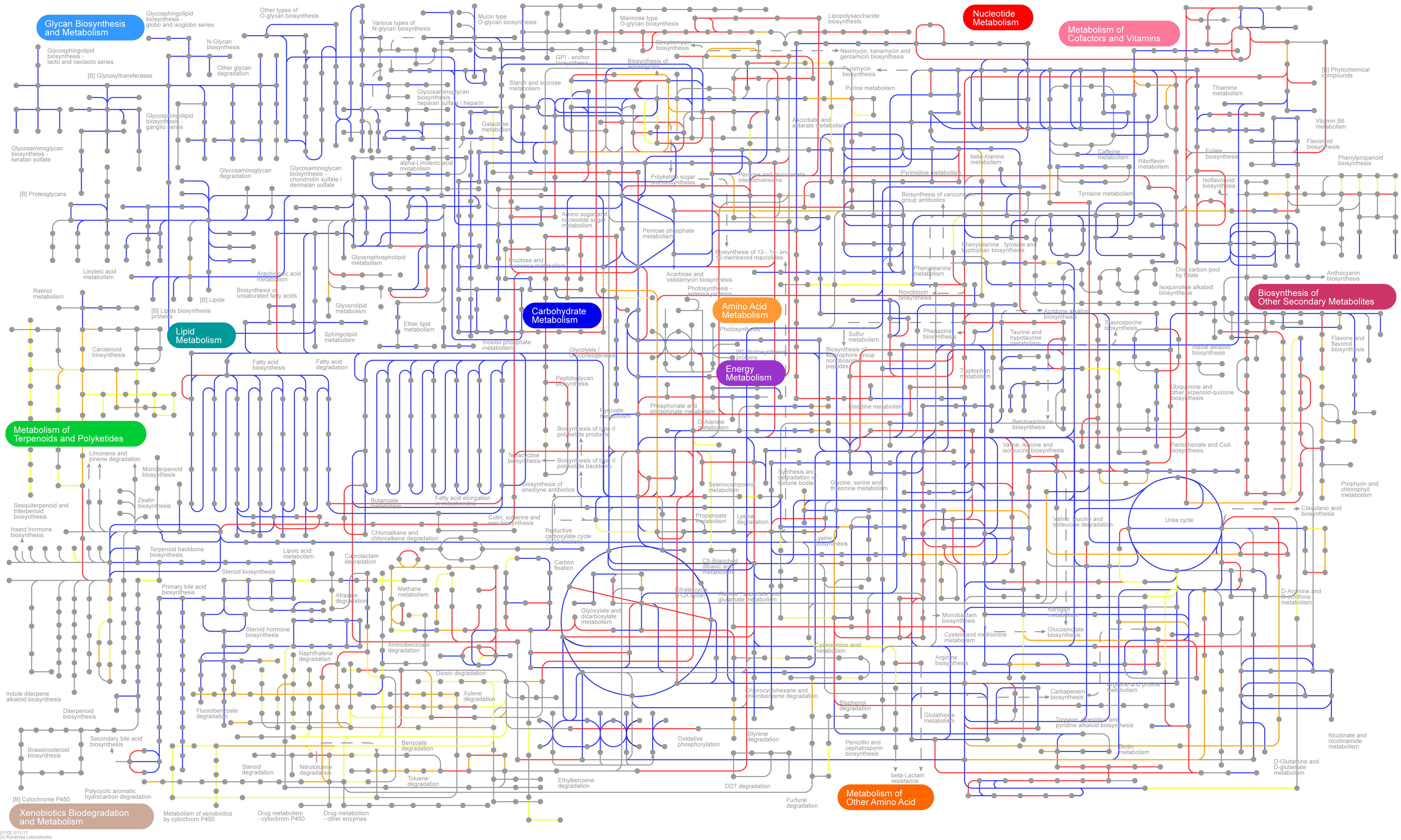

### Supplementary Figure 4

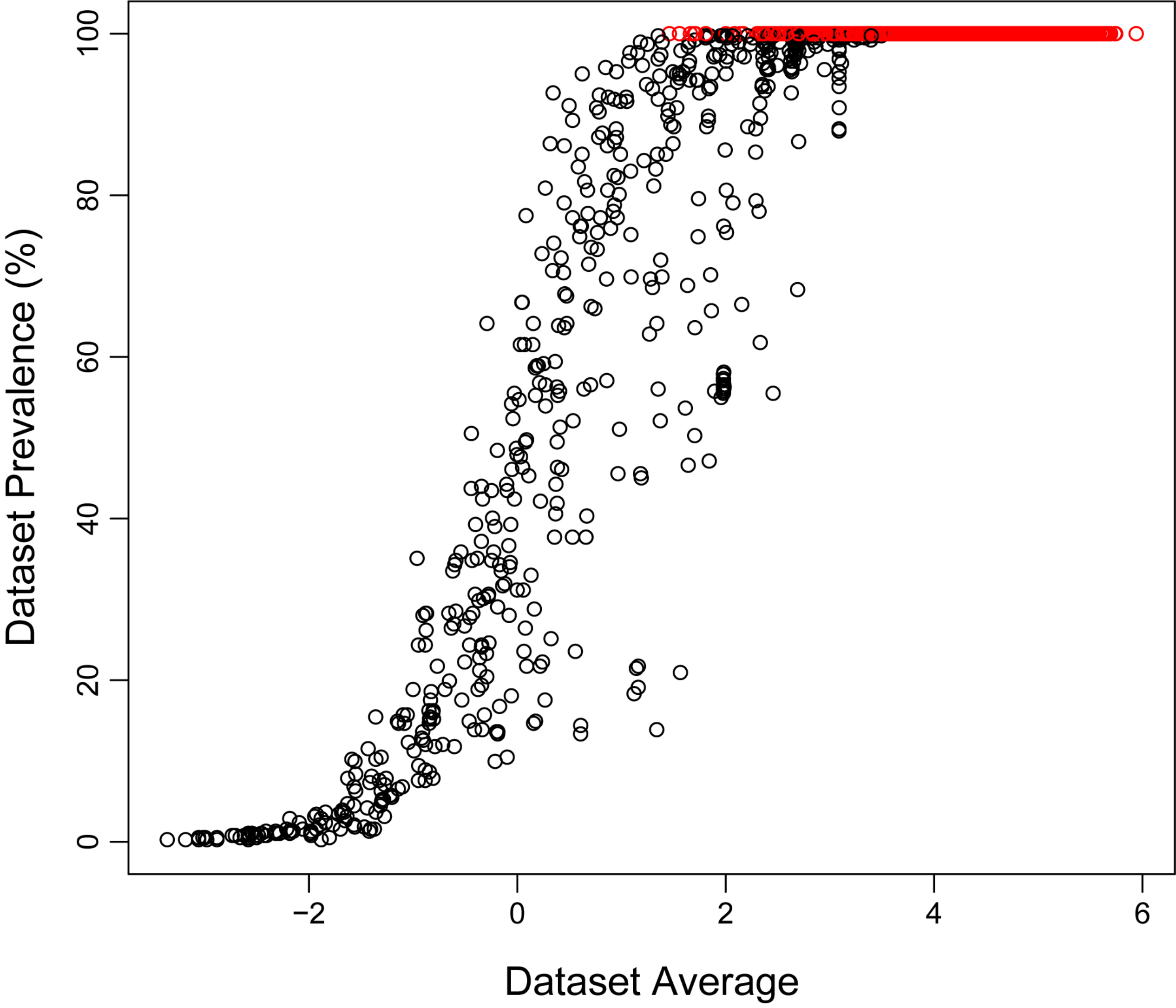
