## Supplemental Data 4 for "A comprehensive human minimal gut metagenome extends the host’s metabolic potential"

**#QIIME Scripts employed**

identify_chimeric_seqs.py -i file.fa -m usearch61 -o usearch_chimeras/ --suppress_usearch61_ref

filter_fasta.py -f file.fa -o fileCC.fa -s usearch_chimeras/chimeras.txt –n

pick_otus.py -i fileCC.fa -o picked/ -m usearch61_ref -C

pick_rep_set.py -i fileCC_otus.txt -f fileCC.fa -o rep_set.fasta

assign_taxonomy.py -i rep_set.fasta -o rep_set.assignedTax -m rdp

make_otu_table.py -i fileCC_otus.txt -t rep_set.assignedTax/rep_set_tax_assignments.txt -o table.biom

**#Common (minimum) depth employed for each dataset**

*Flemish*: 8,383

*TwinsUK*: 14,082

*Global*: 319,000
